# Validation of the protein kinase *PfCLK3* as a multi-stage cross species malarial drug target

**DOI:** 10.1101/404459

**Authors:** Mahmood M Alam, Ana Sanchez-Azqueta, Omar Janha, Erika L. Flannery, Amit Mahindra, Kopano Mapesa, Nicolas Brancucci, Yevgeniya Antonova-Koch, Kathryn Crouch, Nelson Victor Simwela, Jude Akinwale, Deborah Mitcheson, Lev Solyakov, Kate Dudek, Carolyn Jones, Cleofé Zapatero, Christian Doerig, Davis C. Nwakanma, Maria Jesús Vázquez, Gonzalo Colmenarejo, Maria Jesús Lafuente, Maria Luisa Leon, Andrew P. Waters, Andrew G. Jamieson, León Elena Fernandez Alvaro, Matthias Marti, Elizabeth A. Winzeler, Francisco Javier Gamo, Andrew B. Tobin

**Author notes:** Contributed equally to this work (Mahmood M Alam, Ana Sanchez-Azqueta).

## Abstract

The requirement for next generation anti-malarials to be both curative and transmission blockers necessitate the identification of molecular pathways essential for viability of both asexual and sexual parasite life stages. Here we identify a selective inhibitor to the *Plasmodium falciparum* protein kinase *Pf*CLK3 which we use in combination with chemogenetics, whole genome sequencing and transcriptomics to validate *Pf*CLK3 as a druggable target acting at multiple parasite life stages. Consistent with the proposed role of *Pf*CLK3 as a regulator of RNA splicing, inhibition results in the down-regulation of >400 genes essential for parasite survival. Through this mechanism, blocking *Pf*CLK3 activity not only results in rapid killing of asexual blood stage parasites but is also effective on sporozoites and gametocytes as well as showing parasiticidal activity in all *Plasmodium* species tested. Hence, our data establishes *Pf*CLK3 as a target with the potential to deliver both symptomatic treatment and transmission blocking in malaria.

## Introduction

Despite artemisinin-based combinations therapies offering effective frontline treatment for malaria there are still over 200 million cases of malaria worldwide per annum resulting in an estimated 0.5 million deaths. This, combined with the fact that there is now clear evidence for the emergence of resistance to not only artemisinin(*1, 2*) but also to partner drugs including piperaquine and mefloquine(*3, 4*) means that there is an urgent need for novel therapeutic strategies to not only cure malaria but also to prevent disease transmission. Global phospho-proteomic studies on the most virulent species of human malaria, *P. falciparum* have established protein phosphorylation as a key regulator of a wide range of essential parasite processes (*5-8*). Furthermore, of the 65 eukaryotic protein kinases in the parasite kinome(*9*), over half have been reported to be essential for blood stage survival(*8, 10-12*). These studies, together with the generally accepted potential of targeting protein kinases in the treatment of numerous human diseases(*13, 14*), suggests that inhibition of parasite protein kinases might offer a viable strategy for the treatment of malaria(*6, 15*). To directly test this hypothesis we focused here on one of the four members of the *P. falciparum* cyclin-dependent like ((CLK) protein kinase family, *Pf*CLK3 (PF3D7_1114700), an essential protein kinase in maintaining the asexual blood stage of both *P. falciparum*(*8*) and *P. berghei* (*10*). In mammalian cells the CLK protein kinase family and the closely-related SRPK family are crucial mediators of multiple phosphorylation events on splicing factors, including serine-arginine-rich (SR) proteins, which are necessary for the correct assembly and catalytic activity of spliceosomes (reviewed in REF 15)(*16*). A key member of the human CLK-family is the splicing factor kinase PrP4 kinase, which homology-based studies have identified as the closest related human kinase to *Pf*CLK3(*17, 18*). Prp4 kinase plays an essential role in the regulation of splicing by phosphorylation of accessory proteins associated with the spliceosome complex(*19*). The finding that *Pf*CLK3 can phosphorylate SR proteins *in vitro*(*20*) supports the notion that *Pf*CLK3, like the other members of the *Pf*CLK family(*17*), plays an essential role in parasite pre-mRNA processing(*18*).

### High throughput screen identifies selective PfCLK3 inhibitors

We established high-throughput inhibition assays for two essential members of the PfCLK family namely *Pf*CLK1 and *Pf*CLK3. Both of these protein kinases were purified as active recombinant proteins **(Supplementary Figure S1a)** and used in a high throughput time-resolved florescence resonance energy transfer (TR-FRET) assay optimised for 1536-well format using CREBtide and MBP-peptide as substrates for *Pf*CLK1 and *Pf*CLK3, respectively **(Supplementary Figure S1b-g).** In these assays *Pf*CLK3 and *Pf*CLK1 showed Km values for ATP of 10 µM and 30 µM, linear increase in activity associated with protein concentration between a range of 5-30 nM kinase concentration, and stabilities of signal over the assay period (e.g. >60 minutes) **(Supplementary Figure S1b-g)** as well as robust reproducibility with Z’ values >0.7. Using these conditions **(Supplementary Figure S1h)** 24,619 compounds, comprised of 13,533 compounds in the Tres Cantos Anti-Malarial Set (TCAMS)(*21*), 1115 in the Protein Kinase Inhibitor Set (PKIS)(*22*) and 9,970 MRCT index library(https://www.lifearc.org/mrc-technology-launches-small-molecule-compound-library-access-scheme/), were screened for inhibition of *Pf*CLK1 and *Pf*CLK3 at a single dose (10μM). Hits were defined as those compounds that were positioned >3 standard deviations from the mean of the % inhibition distribution curve **(Figure 1a,b)** and which also showed >40% inhibition. This identified 2579 compounds (consisting of MRCT=250, PKIS=4, TCAMS=2325) which together with the 259 compounds identified as *“the kinase inhibitor set*” in the Medicines for Malaria Venture (MMV) box(*23*) were used to generate concentration inhibition curves **(Figure 1c) (Supplementary Table S1)**. Based on the selectivity criteria of more than a >1.5 log fold difference in pIC_50_, 28% of the hits showed specific inhibition of *Pf*CLK1 and 13% specifically inhibited *Pf*CLK3 whilst 23% of the compounds inhibited both *Pf*CLK3 and *Pf*CLK1 with the remainder (36%) being inactive **(Figure 1c,d) (Supplementary Table S1).** Exemplar molecules from each of these three classes are shown in **Figure 1e**. In the case of the *Pf*CLK3 selective molecules these fell into 19 clusters and 42 singletons. Highlighted in Figure 1c, is TCMDC-135051 which showed the highest selectivity and efficacy for inhibition of *Pf*CLK3. Furthermore, TCMDC-135051 showed ˜100 fold lower activity against the closely related human kinase hCLK2 (29% sequence identity with *Pf*CLK3) **(Supplementary Table S1)**. TCMDC-135051 was a member of a series of molecules with the same chemical scaffold that all showed similar inhibitor activity against PfCLK3 **(Supplementary Figure S2a,b,c).** Note that the TCMDC-135051 is part of the TCAMs and has previously been shown to have anti-parasiticidal activity (EC_50_=320nM) and the structure previously been published (*21*). Resynthesis of TCMDC-135051 together with NMR analysis has however determined the correct structure for TCMDC-135051 to be the one shown in Figure 1e.

**Figure 1.**
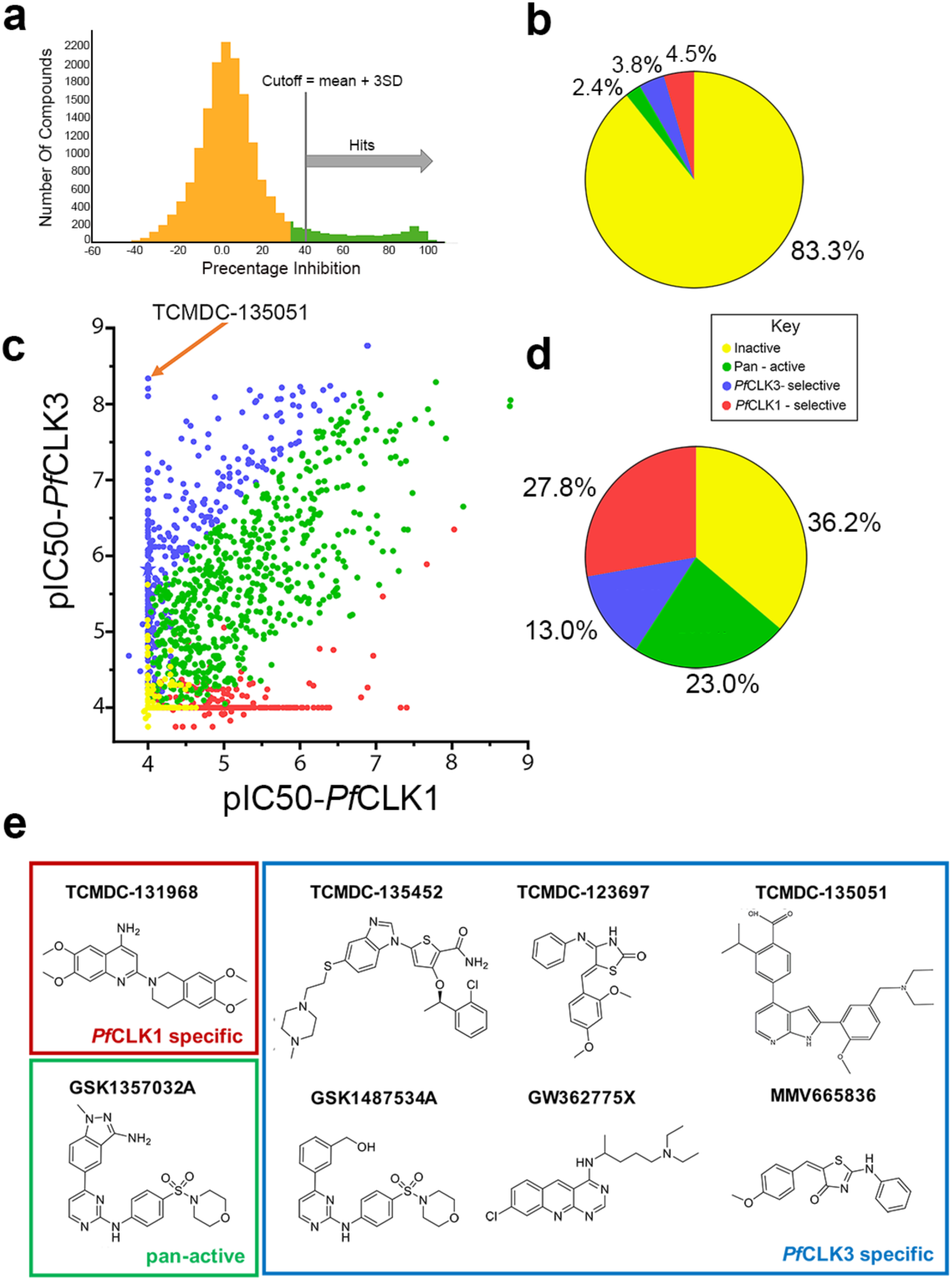
High-throughput Screen Identifies Inhibitors to *Pf*CLK1 and *Pf*CLK3. **(a)** Percent Inhibition distribution pattern of compounds screened against *Pf*CLK3, binned in 5% intervals. Active “hit’ compounds were defined as those that were positioned > 3SDs from the mean. **(b)** Pie-chart summary of the primary single dose screen. **(c)** Hit compounds were used in concentration response curves. Shown is a comparison of pIC_50_ values for inhibition of *Pf*CLK3 vs *Pf*CLK1. TCMDC-135051 is highlighted as the most potent and selective *Pf*CLK3 hit. **(d)** The same data as shown in c but in pie chart format of compounds designated as inactive, pan-active (activity against both *Pf*CLK1 and *Pf*CLK3), and those selective for either *Pf*CLK1 or *Pf*CLK3. **(e)** Chemical structures of eight exemplar *Pf*CLK1 and *Pf*CLK3 compounds that fell into one of three categories, *Pf*CLK1 specific, *Pf*CLK3 specific and *Pf*CLK1/*Pf*CLK3 active (pan active).

In gel-based counter screens against the *P. falciparum* protein kinases, *Pf*PKG and *Pf*CDPK1 **(Supplementary Figures S3a,b,)** compound TCMDC-135051 showed no significant inhibitory activity at concentrations of 2μM, a concentration where TCMDC-135051 completely inhibited *Pf*CLK3 activity **(Supplementary Figures S3c)**.

### Parasite strains resistant to TCMDC-135051 show mutations in *Pf*CLK3

We next sought to confirm that *Pf*CLK3 was the target of TCMDC-135051 *in vivo* activity. Exposing *P. falciparum* Dd2 parasites to increasing concentrations of TCMDC-135051 **(Figure 2a),** resulted in emergence of three independent lines that showed decreased sensitivity to TCMDC-135051 but no change in sensitivity to chloroquine or artemisinin **(Figure 2b, Table 1)**. Whole genome sequencing of the three resistant lines revealed mutations in *Pf*CLK3 (lines TM051A and TM051C) and a mutation in the putative RNA processing protein, PfUSP-(PF3D7_1317000)(line TM051B, **Table 2)**. The resistant clone TM051A showed the smallest change in sensitivity to TCMDC-135051 (4.2 fold shift in the pEC_50_ compared to parental Dd2 parasites). Likewise, examination of the *in vitro* enzymatic properties of mutant *Pf*CLK3 containing the P196R mutation found in TM051A did not detect any significant changes in enzyme kinetics or sensitivity to inhibition by TCMDC-135051 compared to the wild type kinase suggesting that this mutation could potentially stabilise the protein or be otherwise involved in the regulation of the interaction between *Pf*CLK3 and other proteins within the parasite. The line, TM051C, showed the largest degree of resistance to TCMDC-135051 with a shift in the EC_50_ of >11 fold in the death curve **(Figure 2c, Table 1)**. Evaluation of the enzymatic properties of mutant *Pf*CLK3 containing the H259P mutation found in TM051C revealed that the mutant kinase possessed ˜3 times the activity of the wild type kinase whereas the Km for ATP was similar between mutant and wild type kinases **(Figure 2d,e)**. Combined these findings support the notion that the parasiticidal activity of TCMDC-135051 is via inhibition of *Pf*CLK3.

**Table 1:** Adaptive resistance to TCMDC-135951. Dd2 parasites were exposed to TMDC-135051 and three lines isolated that were less sensitive to TCMDC-135051 but with unchanged sensitivity against artemisinin (ART) and chloroquine (CQ). Shown are the IC_50_ values associated with each line and the fold change compared to Dd2 parent parasites. Data is the mean ± S.E.M of three experiments.

| Line | ART<br>(μM) | CQ<br>(μM) | TCMDC-135051<br>(μM) | Fold Change |
| --- | --- | --- | --- | --- |
| Dd2-parent | 0.028 ± 0.006 | 0.169 ± 0.059 | 0.113 ± 0.021 |  |
| TM051A | 0.022 ± 0.006 | 0.142 ± 0.034 | 0.471 ± 0.007 | 4.2 |
| TM051B | 0.018 ± 0.004 | 0.263 ± 0.096 | 0.613 ± 0.044 | 5.4 |
| TM051C | 0.023 ± 0.010 | 0.180 ± 0.054 | 1.25 ± 0.217 | 11.1 |

**Table 2.**
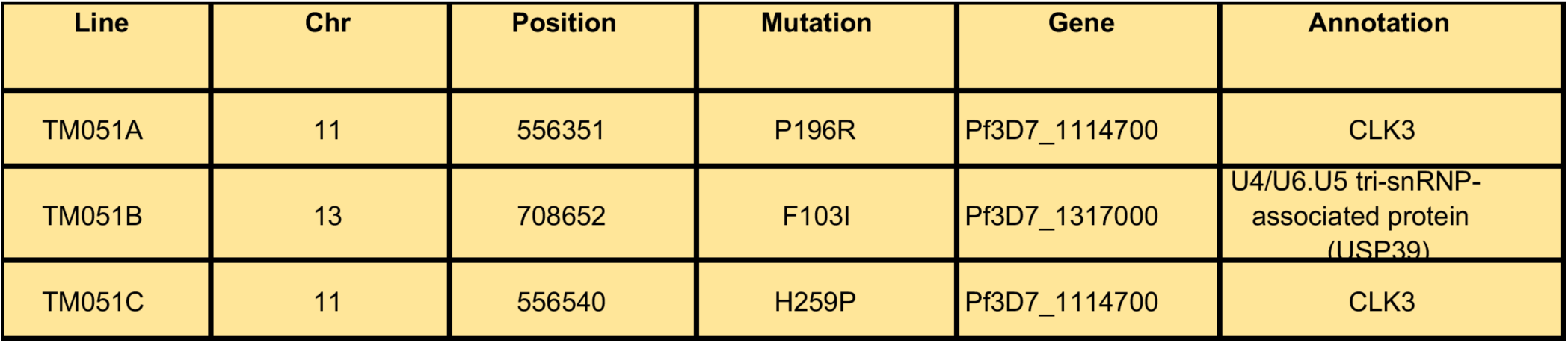
Mutations associated with adaptive resistance to TCMDC-135051. Genomic DNA was isolated from each of three lines derived from adaptive resistance to TMDC-135051 and whole genome sequencing was performed. DNA libraries were prepared using a Nextera-XT kit and sequenced on the Illumina Mi-Seq with 250 bp paired-end reads. Average coverage for each strain was 29.9x. Shown are the single nucleotide variants identified as unique compared to the parent Dd2 line.

**Figure 2.**
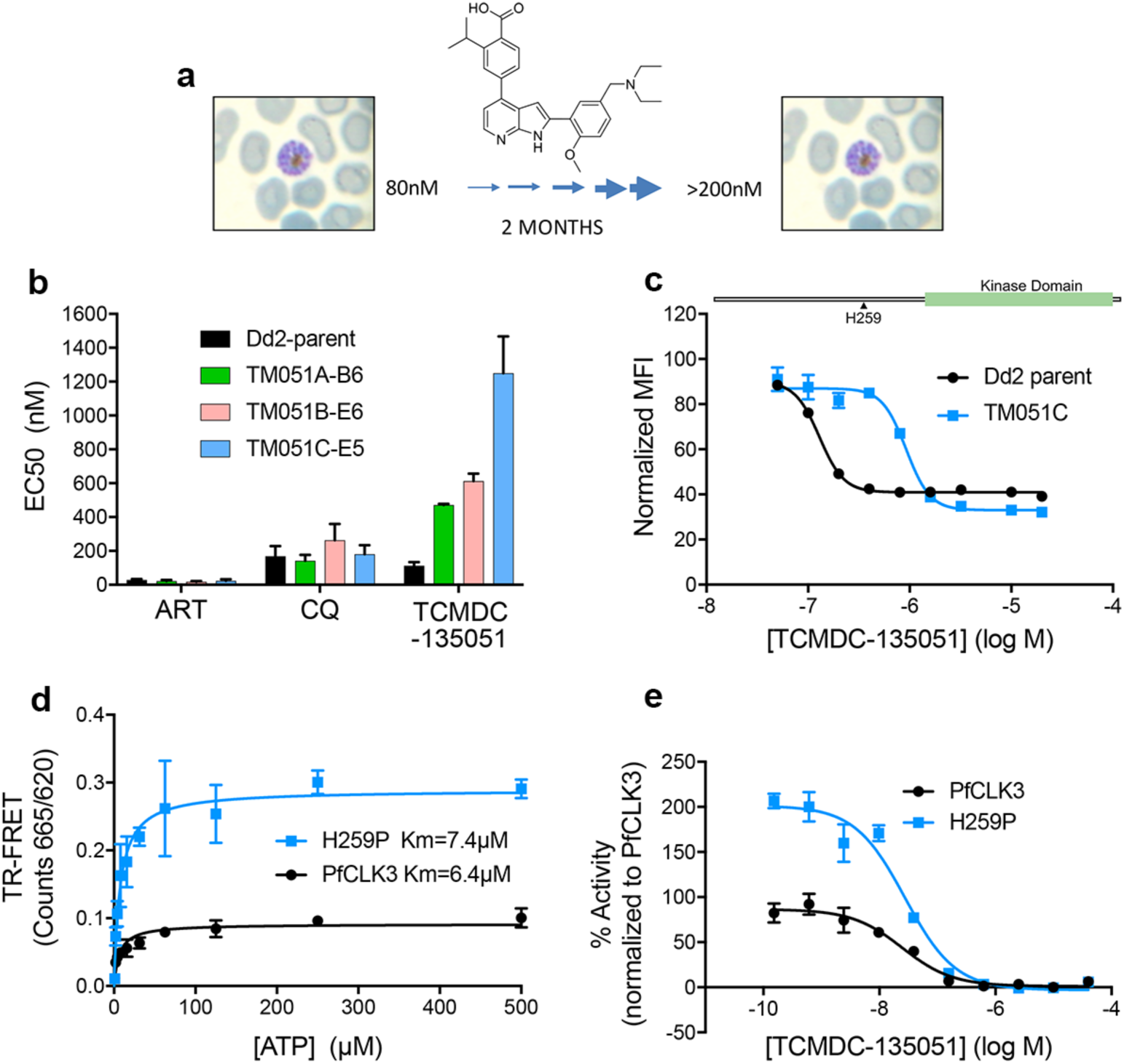
Parasites adapted to become less sensitive to TCMDC-135051 harboured mutations in *Pf*CLK3. **(a)** Schematic of the generation of TCMDC-135051 resistant lines. Over a 2 month period Dd2 parasites were cultured with increasing concentrations of TCMDC-135051. **(b)** Three lines were isolated that were less sensitive to TCMDC-135051 but with unchanged sensitivity against artemisinin (ART) and chloroquine (CQ). **(c)** The line showing the greatest change in sensitivity to TCMDC-135051, TM051C, expressed a mutant form of *Pf*CLK3 where histidine 259 was substituted by a proline (illustrated). Shown are death curves for parental and TM051C lines. **(d)** Enzyme activity of recombinant *Pf*CLK3 and the H259P mutant were determine at varying ATP concentrations to derive a Km for ATP. **(e)** TCMDC-135051 kinase inhibition curves for *Pf*CLK3 and the H259P mutant at Km ATP concentrations for each enzyme. Data presented is the mean ± S.E.M of at least 3 independent experiments.

### Genetic target validation of *Pf*CLK3

To further validate *Pf*CLK3 as the target for parasiticidal activity of TCMDC-135051 a genetic approach was taken in which a recombinant *Pf*CLK3 variant was designed that showed reduced sensitivity to TCMDC-135051. To generate this variant, advantage was taken of the highly selective inhibition of *Pf*CLK3 over *Pf*CLK1 shown by TCMDC-135051. This selectivity was observed despite these two kinases showing 31% identity in primary amino acid sequence. By exchanging amino acids within the sub-domain IV of the *Pf*CLK3 kinase domain with equivalent residues in the kinase domain of *Pf*CLK1 a variant of *Pf*CLK3 was generated where glycine 449 in *Pf*CLK3 was substituted with a proline residue (G449P) **(Figure 3a)**. This recombinant variant protein showed a ˜3 fold log shift in sensitivity for inhibition by TCMDC-135051 **(Figure 3b,c)** (*Pf*CLK3 pIC_50_=7.35±0.12, G449P pIC_50_=4.66±0.16). The G449P variant als0 showed a slightly lower enzymatic activity **(Figure 3d)** (*Pf*CLK3 Vmax=1.24, G449P Vmax=0.88) but higher Km for ATP **(Supplementary Figure S4)** (*Pf*CLK3 Km=6.29, G449P Km=81.3) compared to wild type *Pf*CLK3. Single cross over homologous recombination targeting the *Pf*CLK3 locus with a construct designed to insert the coding sequence for G449P mutant **(Figure 3e)** generated two independent clones (A3 and A8), which expressed the G449P mutant in place of the wild type *Pf*CLK3 **(Figure 3f,g)**. Integration of the plasmid at the target locus was verified by PCR of genomic DNA **(Figure 3f)** and western blotting confirmed expression of the G449P mutant which was epitope tagged with a haemaggultinin (HA) tag at the C-terminus **(Figure 3g).** The activity of TCMDC-135051 in parasite viability assays was seen to be significantly reduced by ˜1.5 log units in both clones of G449P **(Figure 3h)** (pEC_50_ for TCMDC-135051 in the Dd2 wild type = 6.35 ± 0.038 and in the A3 strain = 4.86 ± 0.13 and the A8 strain = 4.94 ± 0.051), providing further evidence that TCMDC-135051 kills parasites via inhibition of *Pf*CLK3.

**Figure 3.**
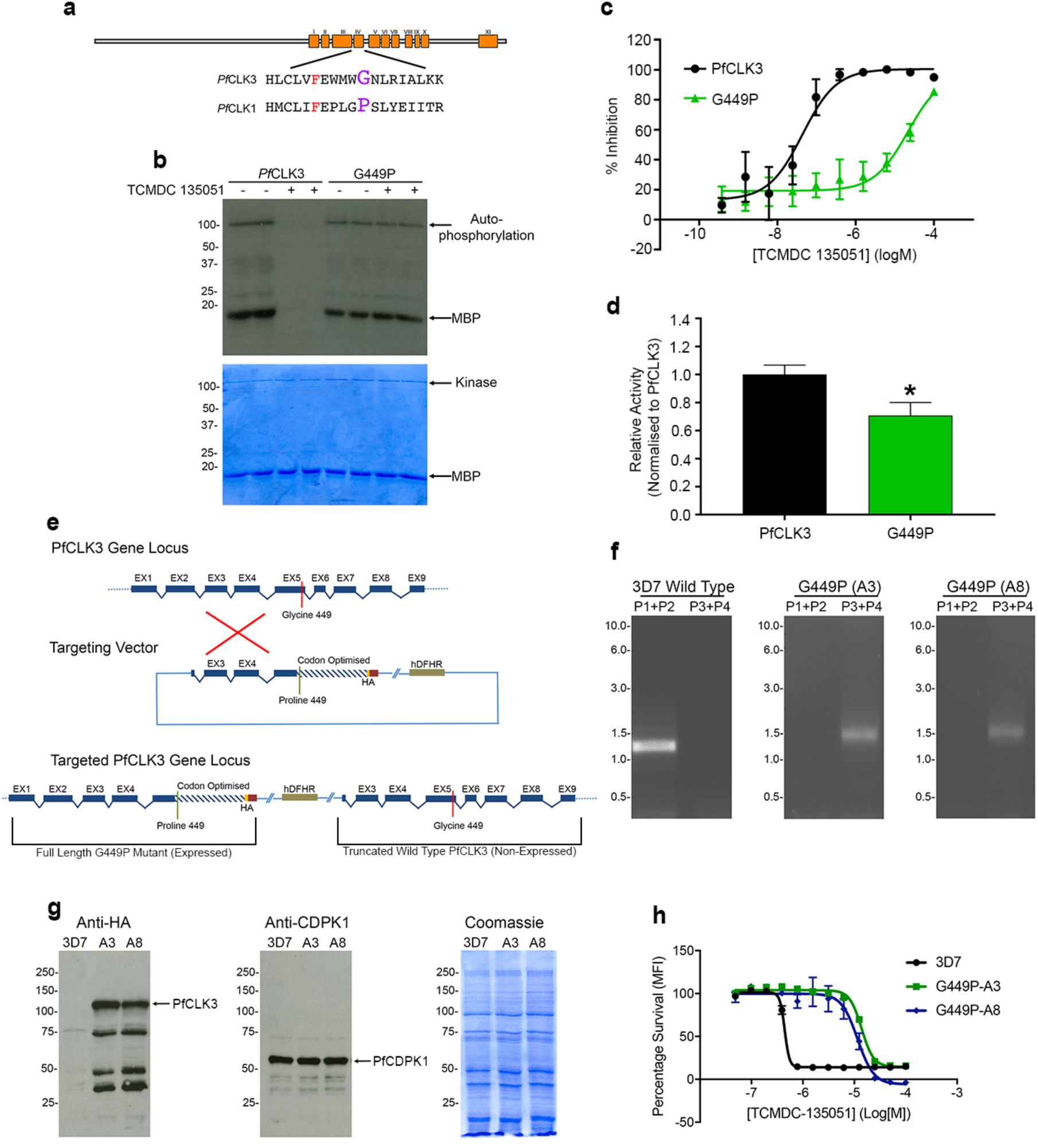
Chemogenetic validation of *Pf*CLK3 as a target for the parasiticidal activity of TCMDC-135051. **(a)** Schematic of the primary amino sequence of *Pf*CLK3 showing the 11 kinase subdomains and the amino acid sequence of subdomain IV of *Pf*CLK1 and *Pf*CLK3. **(b)** Gel based assay of the phosphorylation of myelin basic protein (MBP) by *Pf*CLK3 and a variant where glycine at position 449 was substituted for a proline (G449P). The top gel is an autoradiograph and the bottom a Commassie stain of the same gel which serves as a loading control. The position of MBP and auto-phosphoryated kinase is shown. **(c)** Concentration kinase inhibition curves of TCMDC-135051 inhibition of *Pf*CLK3 and the G449P mutant. **(d)** Maximal kinase activity of recombinant *Pf*CLK3 compared to the activity of the G449P mutant. **(e)** Schematic of gene targeting strategy that would result in the expression of the G449 mutant (containing a triple HA-tag) in place of wild type *Pf*CLK3. **(f)** The recombination event illustrated in “e” was identified in cloned G449P parasite cultures by PCR (A3 and A8). **(g)** Expression of the triple HA-tagged G449P mutant in genetically engineered parasite cultures (G449P) was determined by Western blotting. Gel on the left probes lysates with anti-HA antibodies, the gel in the middle is a loading control probed with anti-CDPK1 antibodies and the gel on the right is a Commassie stain of the lysate preparations used in the Western blots. **(h)** The growth inhibition curves of TCMDC-135051 against parent 3D7 parasites and G449P parasites (A3 and A8). The data shown in the graphs are the mean ±S.E.M of at least three independent experiments. *p<0.05 t-test.

### Inhibition of *Pf*CLK3 prevents trophozoite to schizont transition

To characterize the phenotypic response to *Pf*CLK3 inhibition and to understand *PfCKL3* function, *P. falciparum* 3D7 parasites synchronised at ring stage (time point zero) were treated with TCMDC-135051 (1μM). The parasites progressed to late ring stage (time point 20 hours) **(Figure 4a)** but did not progress further to trophozoite stage (time point 30 and 40 hours), arresting with a condensed and shrunken appearance **(Figure 4a)**. Similar effects were observed if the parasites were treated at mid ring stage (time point 10 hours). Treatment of parasite at later time points (20 hours or 30 hours) blocked development of the parasite from trophozoite to schizont stage and led to condensed staining of nuclei, which is typical of stressed and non-viable parasites. The fact that the parasites at the schizont stage were not viable was further evidenced by the absence of ring stage parasites when the culture was allowed to continue to the 50-hour time point **(Figure 4a)**. These data indicated that *Pf*CLK3 inhibition prevented the transition of the parasites at early (ring to trophozoite) as well as late stages (trophozoite to schizont) of development and importantly did not allow parasites to reach to the next invasion cycle **(Figure 4a)**. These data further indicated that *Pf*CLK3 inhibition resulted in rapid killing with no evidence that the compound results in quiescence from which the parasite can recover following drug withdrawal. These features were confirmed in parasite reduction rate (PRR) assays, which showed that treatment of parasites with 10X pEC_50_ of TCMDC-135051 completely kills the parasite in 48 hours and viable parasites could not be observed despite maintaining the parasite culture for 28 days following withdrawal of TCMDC-135051 **(Figure 4b).** Furthermore, the killing profile is consistent with TCMDC-135051 acting against trophozoite stages, as the PRR assay started with ring-rich stage parasites (24 h lag phase to reach maximal killing rate and full parasite eradication afterwards(*24*)).

**Figure 4.**
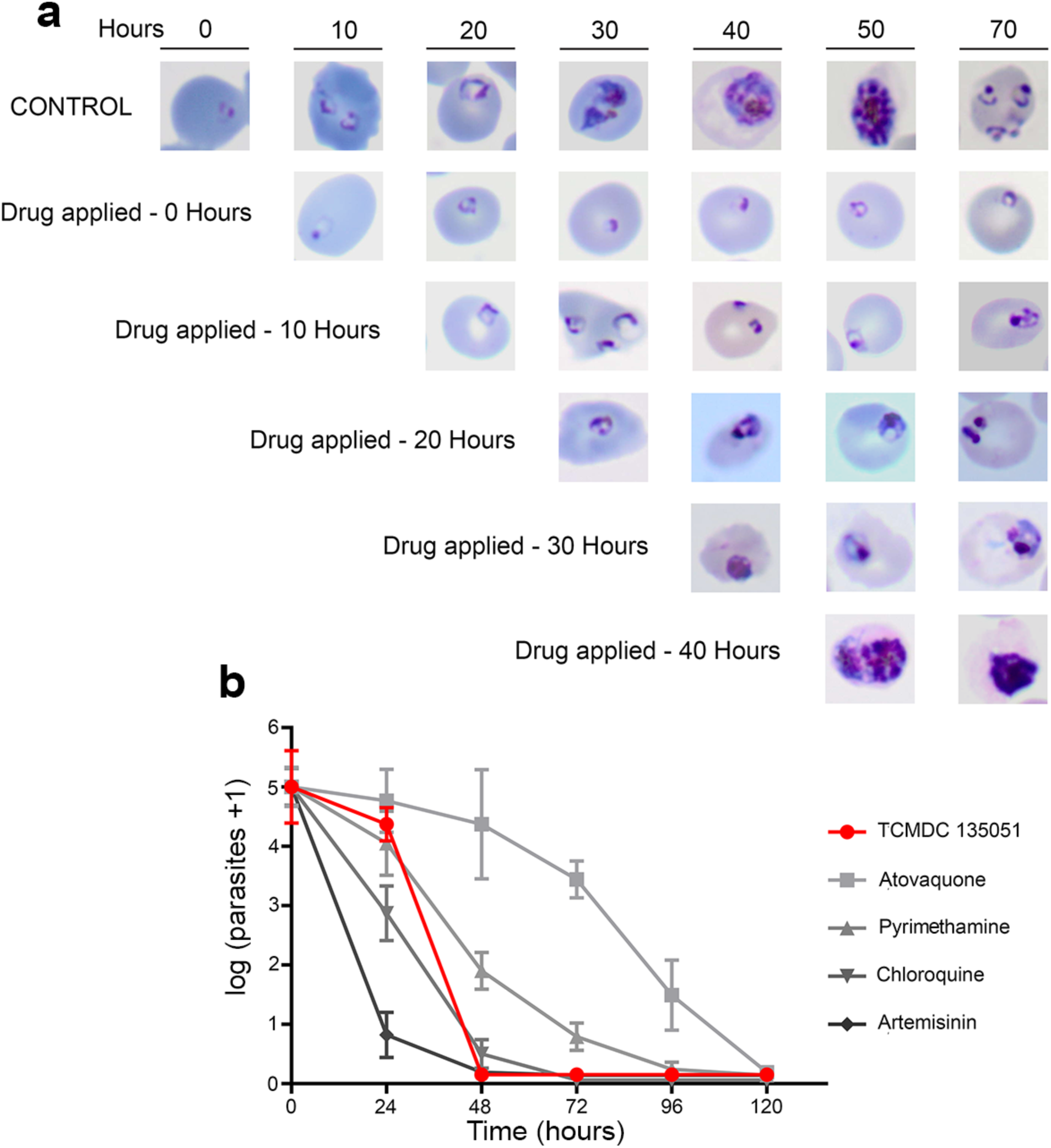
Inhibition of *Pf*CLK3 prevents trophozoite to schizont transient and kills the parasite with rapid kinetics. **(a)** Smears of synchronised blood stage *P. falciparum* cultures following treatment with TCMDC-135051 (2μM) were taken at the indicated times after TCMDC-135051 administration. **(b)** The *in vitro* parasite reduction rate assay was used to determine onset of action and rate of killing as previously described(*24*). *P. falciparum* was exposed to TCMDC-135051 at a concentration corresponding to 10 × pEC_50_. During the experiment, medium was exchanged and drugs replaced every 24 hours. Aliquots corresponding to 10^5^ parasites were taken out at defined points, washed free of drug and the parasites diluted by serial dilution and cultured in fresh medium and with fresh erythrocytes. Parasite growth was subsequently monitored after 28 days and the number of parasites in the culture before drug wash out back-calculated. Four independent serial dilutions were done with each sample to correct for experimental variation. The data presented represents the mean ±S.E.M. Previous results reported on standard anti-malarials tested at 10 × pEC_50_ using the same conditions are shown for comparison(*24*).

### Inhibition of *Pf*CLK3 disrupts transcription

The human kinases, PrP4 and hCLK2, are closely related to *Pf*CLK3 and are protein kinases involved in the regulation of RNA splicing, primarily through the phosphorylation of serine/arginine rich splicing proteins (SR proteins) and other accessory proteins of the spliceosome complex(*19*). Our earlier studies demonstrated that *Pf*CLK3 similarly phosphorylate parasite SR-proteins PfSFRS4 and PfSRSF12(*20*) and the fact that PfSRSF12 is involved in RNA processing in *P. falciparum*(*25*) supports the notion that *Pf*CLK3 is a key regulator of RNA processing. Interestingly, one of the resistant strains, TM051B, emerging from the adaptive-resistance experiments contained a mutation in the RNA processing protein, PfUSP-39, a protein known to be essential for spliceosome complex formation (*26*). As such to investigate the parasiticidal mode of *Pf*CLK3 inhibition, changes in gene transcription in parent Dd2 parasites and drug-resistant stain TM051C, in response to exposure to TCMDC-135051 was evaluated. RNA isolated from trophozoite stage parasites was extracted following a short 60 minute treatment with TCMDC-135051 (1μM), a period of time in which the Dd2 and TM051C parasites maintained normal histology. Genome-wide transcriptional patterns were determined using oligonucleotide microarray chips that probed 5752 *P. falciparum* genes (*27*). Under these conditions 779 gene transcripts were significantly down-regulated in response to *Pf*CLK3 inhibition in the Dd2 parasites and 155 genes were up-regulated **(Figure 5a, Supplementary Table S2)**. That the majority of these transcriptional changes were due to inhibition of *Pf*CLK3 and not off-target events was supported by the fact that under the same conditions only 6 genes were up-regulated and 88 down-regulated in the resistant TM051C parasite strain **(Figure 5b, Supplementary Table S3)**. By subtracting the transcriptional changes observed in the TM051C strain, defined here as “off-target”, from those observed with the Dd2 parent, the transcriptional changes due to “on-target” inhibition of *Pf*CLK3 were defined **(Supplementary Table S4)**. Among these “on-target” down-regulated genes were those involved in key parasite processes such as egress and invasion, cytoadherence, parasite protein export and involvement in sexual stages as well as house-keeping functions including metabolism, RNA processing, lipid modification and mitochondrial function **(Figure 5c, Supplementary Table S4)**. Importantly, of the 696 genes identified as down-regulated by *Pf*CLK3 inhibition (**Supplementary Table S4)**, 425 matched those that have recently been determined to be essential for asexual *Plasmodium falciparum* survival (*12*) **(Supplementary Table S4).**

**Figure 5.**
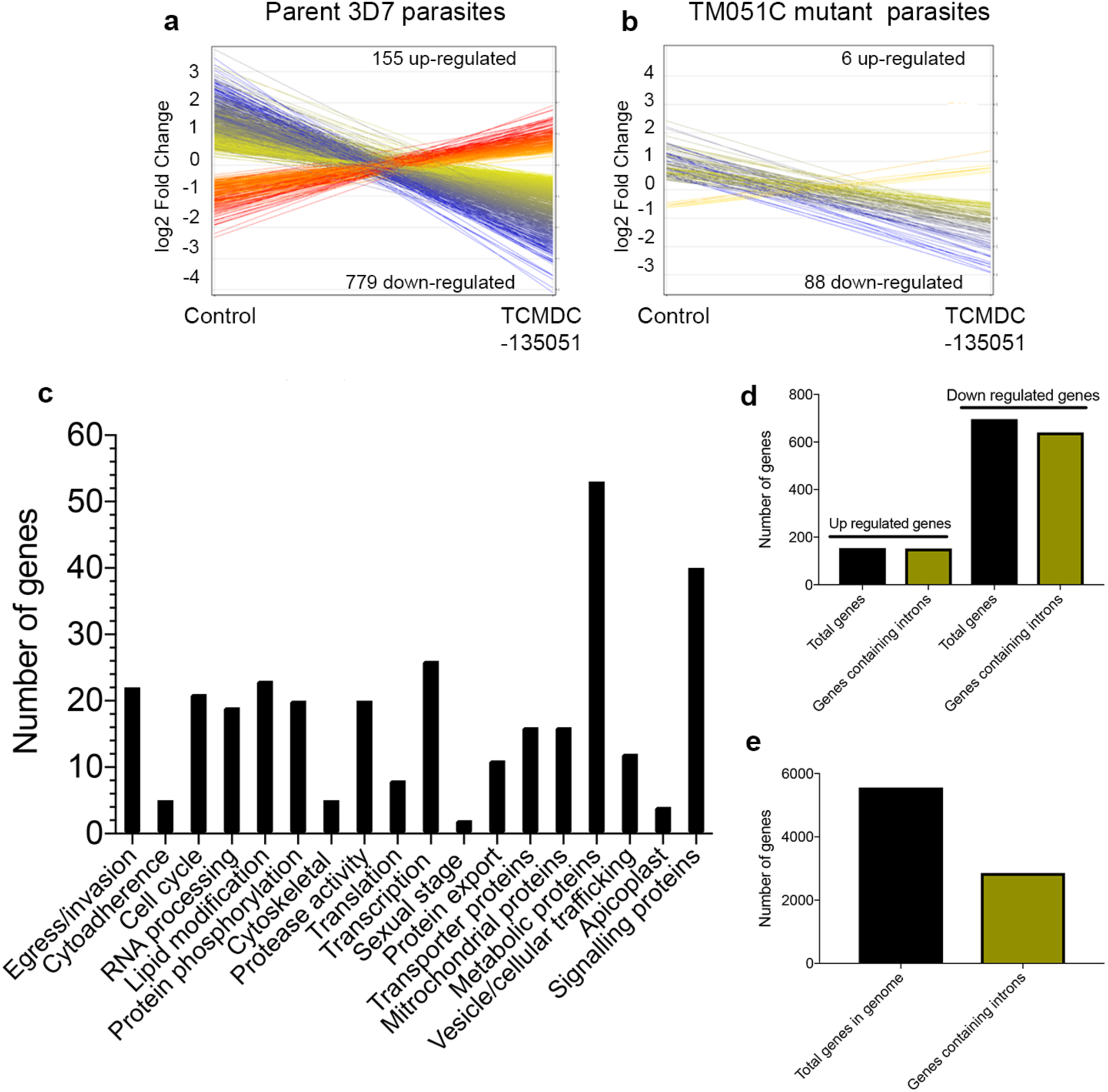
Gene transcription changes following inhibiton of PfCLK3. Illustration of the genes that are designated as significantly changing (Moderate t test n=4) in transcription following treatment with TCMDC-135051 (1μM, 60 mins) of either **(a)** parent Dd2 parasites or **(b)** TM051C mutant parasites. Each line represents the log2 fold change in the probes used in the micro-array. The number of genes represented by the probes is indicated. **(c)** Summary of the parasite processes associated with the genes where transcription is down-regulated following TCMDC-135051 treatment. **(d)** Assessment of intron containing genes contained among genes that are up-regulated and down-regulated in Dd2 parasites following TCMDC-135051 treatment. **(e)** Assessment of intron containing genes in the plasmodium genome (data derived from Plasmodb).

Also of interest was that 93% of the genes that were seen to change in response to *Pf*CLK3 inhibition contained introns **(Figure 5d, Supplementary Table S4)**. This is compared to ˜50% of genes in the *P. falciparum* genome that are annotated as containing introns(*28*) **(Figure 5e).**

### Multi-stage and cross species activity of TCMDC-135051

The current product profile of candidate anti-malarial compounds in the MMV portfolio includes multi-stage activity that would result in not only clearance of asexual blood stage parasites (curative) but also block malaria transmission (*29*) by targeting sexual stage parasites. TCMDC-135051 was tested in an assay developed using the *P. falciparum Pf*2004 parasite strain that shows high levels of gametocyte production (*30*). Importantly, TCMDC-135051 showed potent parasiticidal activity in asexual stage *Pf*2004 **(Figure 6a)** (pEC_50_ in *Pf*2004 = 6.58 ± 0.01) similar to that seen in 3D7 and Dd2 parasites. In addition, TCMDC-135051 inhibited completely early stage II gametocyte development with a pEC_50_ = 6.04 ± 0.11 **(Figure 6b).**

**Figure 6.**
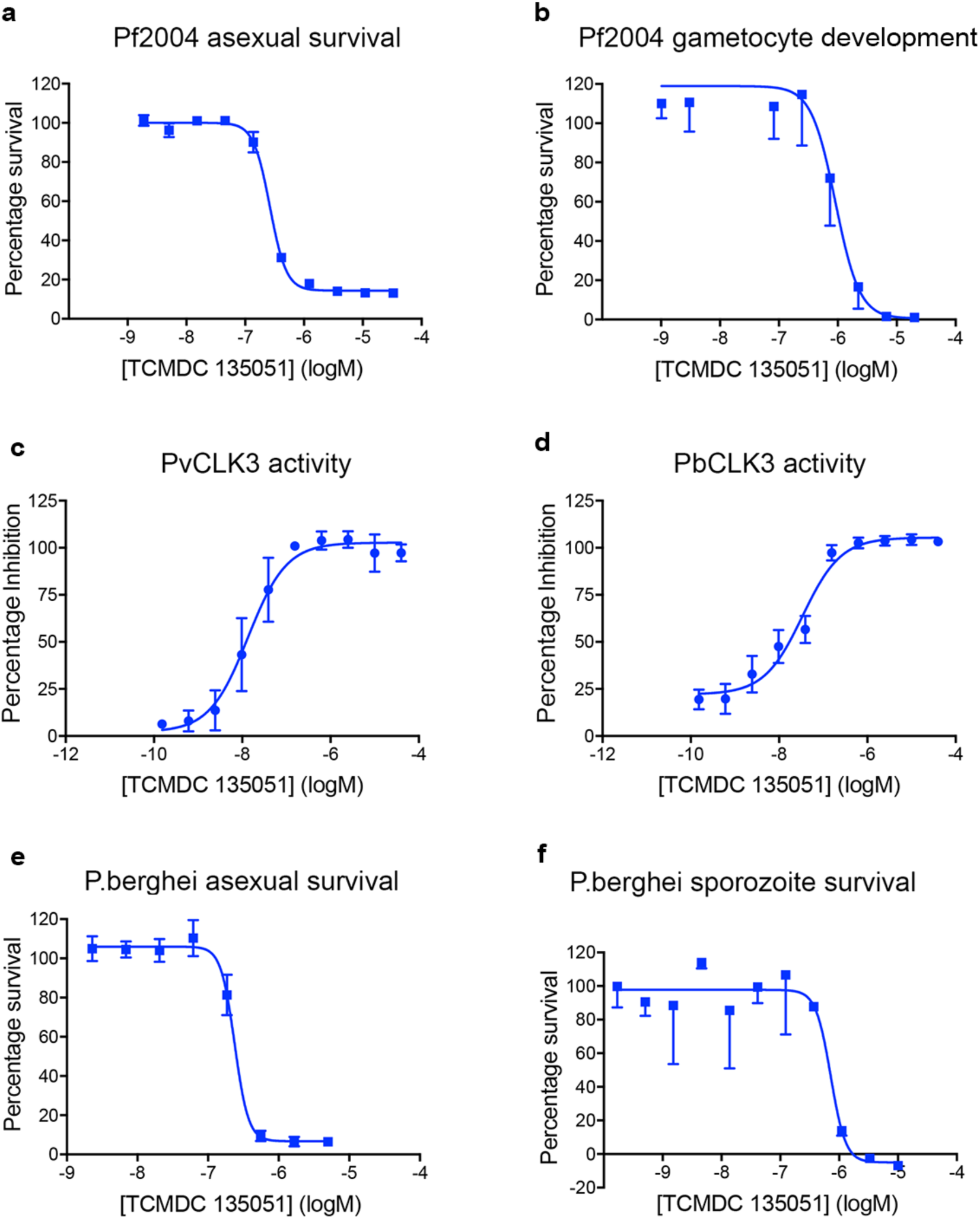
Inhibition of *Pf*CLK3 has parasiticidal activity at multiple parasite stages and on multiple *Plasmodium* species. **(a)** Concentration effect curve of TCMDC-135051 on blood stage *Pf*2004 parasites. **(b)** Activity of TCMDC-135051 on development of gametocytes in *Pf*2004 parasites. **(c)** *In vitro* kinase inhibition concentration curve of TCMDC-135051 against *Pv*CLK3 kinase activity. **(d)** *In vitro* inhibition concentration curve of TCMDC-135051 against *Pb*CLK3 kinase activity. **(e)** Concentration effect curve of TCMDC-135051 on blood stage *P. berghei* parasites. **(f)** Concentration effect curve of TCMDC-135051 on *P. berghei* sporozoite stage parasites. The data shown are the mean ±S.E.M of at least three independent experiments.

Next activity against multiple species of malaria parasite was investigated. It might be predicted that the close similarity between orthologues of CLK3 in different malaria parasites species would result in TCMDC-135051 showing similar activities against CLK3 from different *Plasmodium* species. This indeed was the case as *in vitro* assays using recombinant *Pv*CLK3 (*P. vivax*) and *Pb*CLK3 (*P. berghei*) (**Supplementary Figure S5a,b)**, TCMDC-135051 showed near equipotent inhibition of *Pv*CLK3 and *Pb*CLK3 with pIC_50_ values of 7.86 ± 0.10 and 7.47 ± 0.12 respectively **(Figure 6c and d)**. Based on these findings it can also be predicted that TCMDC-135051 would show activity against the asexual blood stage of *P. berghei* which was found to be the case **(Figure 6e)**. Importantly, TCMDC-135051 also showed potent activity against *P. berghei* sporozoites in a liver invasion and development assay(*31*) in which the compound showed a pEC_50_ value of 6.17 ± 0.10 **(Figure 6f)**, although hepatocyte toxicity was observed but only significantly at 10μM **(Supplementary Figure S6)**. These combined data provide evidence that inhibition of CLK3 had parasiticidal activity in multiple Plasmodium species and at multiple stages that included both asexual (sporozoites and blood stages) and sexual stages (gametocytes).

## Discussion

Here we identify *Pf*CLK3 as a valid and druggable anti-malarial target for both sexual and asexual stages of parasite development. This suggests that targeting *Pf*CLK3 might not only be a novel strategy for developing curative treatments for malaria by clearance of asexual blood stage parasites but the parasiticidal activity afforded by *Pf*CLK3 inhibition at sporozoites and gametocytes would indicate that through this mechanism liver infection and transmission to the insect vector may be affected. Since splicing of essential transcripts occurs at many stages of the parasite life cycle it is attractive to hypothesise that inhibition of *Pf*CLK3, which has been implicated in the phosphorylation of splicing factors necessary for the assembly and activity of the spliceosome (*17, 18, 20*), would have a wide-ranging impact on parasite viability. In support of this notion is the discovery that *Pf*CLK3 inhibition down-regulates over 400 essential parasite transcripts. Interestingly, the majority of down-regulated transcripts are from genes that contain introns (91%), providing further evidence that *Pf*CLK3 is involved in RNA splicing and disruption of this essential process at multiple life cycle stages is the likely mechanism by which inhibitors of *Pf*CLK3 have parasiticidal activity.

The similarity of CLK3 orthologues in *Plasmodium sp*. suggests inhibitors might also have activity across a number of *Plasmodium* species. This was confirmed here by almost equipotent inhibition of the kinase activity of *Pv*CLK3, *Pb*CLK3 and *Pf*CLK3 by TCMDC-135051. This *in vitro* action was mirrored by *in vivo* activity in *P. berghei* and *P. falciparum* asexual blood stage cultures, indicating that inhibition of *Pf*CLK3 might not only satisfy MMV criteria of next generation anti-malarials to show rapid multi-stage parasite killing but also meet the desire for cross species activity (*29*).

One of the major barriers associated with the development of protein kinase inhibitors is the issue of selectivity since the ATP binding pocket, to which the majority of protein kinase inhibitors bind, is very similar between protein kinases(*32*). Here TCMDC-135051 shows surprising selectivity towards *Pf*CLK3 even when compared to its paralogue in *P. falciparum Pf*CLK1 and the closely related human kinase hCLK2. Furthermore, the fact that our transcriptional studies revealed very few off-target events and that adaptive resistance and chemogenetic resistance was associated with single point mutations in *Pf*CLK3 indicates that the selectivity of TCMDC-135051 for *Pf*CLK3 observed *in vitro* was maintained *in vivo*.

Despite abundant evidence that phosphorylation and phospho-signalling is crucial for the viability of both asexual and sexual stages of the malaria parasite (*5, 8, 10, 11*) and reports identifying essential parasite protein kinase targets (*8, 11*), together with the extensive experience of academic and industrial laboratories in the design of protein kinase inhibitor drugs (*14, 32*), the targeting of parasite protein kinases in anti-malarial drug development is only in its infancy (*6, 33*). By focusing on an essential parasite kinase and taking advantage of high throughput phenotypic screens of commercial and academic libraries (*21, 34, 35*) as a starting point to screen for inhibitors, we have identified a probe molecule that has not only established the validity of *Pf*CLK3 as a target in malaria but also determined that this protein kinase is susceptible to selective pharmacological inhibition with small drug-like molecules. In this way, our study lends weight to the argument that targeting the essential parasite protein kinases identified through global genomic studies might be a valid therapeutic strategy in the development molecules that meet many of the criteria set for the next generation of anti-malarial drugs.

## Supplementary Information

### Methods

#### In vitro Kinase Activities and Inhibition Assay

Kinase assays were carried in kinase buffer containing (20 mM HEPES, pH 7.4, 10 mM MgCl_2_, 1 mM DTT, 50 μM adenosine triphosphate (ATP), 0.1 MBq [γ-^32^P]-ATP) using 0.5 μg purified His-tagged recombinant protein kinases. Protein kinases were incubated with exogenous substrate, Histone Type IIA 2 μg; Myelin Basic Protein (MBP) 2 μg; and α-Casein 2 μg; and with a PBS as a negative control for 30 minutes at 37°C.

The reactions were stopped by adding equal volume of 2X Laemmli buffer after incubation and boiled for 2 minutes at 60°C. Samples were separated on 15% SDS-PAGE, stained with Coomassie Brilliant Blue and dried by means of vacuum gel drying. Dried gels were exposed to X-ray film at −80°C and autoradiographs were collected.

For *in vitro* inhibition assays, Compound TCMDC-135051 was added to a final concentration of 2 μM. For control, a no inhibitor reaction tube was setup to exclude any other unknown inhibitory effect. All other reaction conditions remained the same. Quantification of activity was carried out using ImageJ (1.49v).

#### Time Resolve Florescence Energy Transfer (TR-FRET)

Biochemical kinase assays were carried out using time-resolved florescence energy transfer (TR-FRET) to determine kinase activity, Km for ATP and IC50 values for the respective kinases used. TR-FRET reactions were performed using the appropriate amount of kinase (5nM for *Pf*CLK1 and 50 nM for *Pf*CLK3 and G449P) in a kinase buffer (containing 50 mM HEPES, 10 mM MgCl_2_, 2 mM DTT, 0.1% Tween 20, and 1 mM EGTA), U*Light*-labeled peptide substrate (MBP peptide or CREPtide) and an appropriate europium-labeled anti-phospho antibody in two steps.

First, in a 10 μL reaction volume 5 μL of 2X-required enzyme concentration and 5μL of 2X-required substrate mix and cold ATP were incubated in a 384 plate well plate at 37°C for 1 hour. Second, following incubation, adding 30 mM EDTA in 1X Lance detection buffer containing 3nM Europium-labeled anti-phospho specific antibody incubated at room temperature for 1-hour to stop the reaction and enhance detection and this is then read using the ClarioStar.

Kinase substrate phosphorylation results in the Europium-labeled anti-phospho specific antibody recognizing the phosphorylated site on the substrate. The Europium donor fluorophore is excited at 320 or 340 nm and energy is transferred to the U*Light* acceptor dye on the substrate, which finally results in the emission of light at 665 nm. The level of ULight peptide phosphorylation determines the intensity of the emission.

To test for inhibition by small molecules such as TCMDC-135051, serial dilution of the inhibitor (made at 4 times the required concentration) was made and added to the protein mixture before adding the substrate mix at 4X the required concentration. For normalization, a no kinase and a no inhibitor reaction wells were included and all experiments were run in triplicate.

#### Expression and purification of *Pf*CLKs, *Pf*CDPK1 and *Pf*PKG

Gene encoding the kinase domain of *Pf*CLK1 (residues 534-857) was amplified using *P. falciparum* (3D7) genomic DNA and primers CLK1-FamB1 and CLK1-FamB2. The amplified product was cloned in plasmid pLEICS-05 (University of Leicester) as has been previously described for cloning of *Pf*CLK3 (*1*). Plasmids encoding the kinase domain of *Pf*CLK1 (pLEICS-05_*Pf*CLK1 (residues 534-875-3xHIS)) and full length *Pf*CLK3 (pLEICS-05_*Pf*CLK3 (residues-1-669-5xHIS)) were used to transform BL21 Codon Plus (DE3) *Escherichia coli* competent cells (Agilent Technologies). Protein expressions were induced for 4 hours at 37°C for *Pf*CLK3 and at 22°C for *Pf*CLK1 after the addition of 0.1 mM IPTG. Cell pellets were lysed by sonication in Lysis Buffer (20 mM Tris/HCl, pH 8.0, 300 mM NaCl, 1 mM DTT and 10 mM Imidazole) and EDTA-free protease inhibitor cocktail, followed by centrifugation at 8000 g. Clarified supernatants were loaded onto Ni-NTA Superflow cartridges (Qiagen), washed using lysis buffer, and eluted with lysis buffer containing 300mM Imidazole. Protein was dialysed in buffer containing 20 mM Tris/HCl, pH 7.4, 150 mM NaCl, 1 mM MgCl_2_ and 10%Glycerol and aliquots were stored at −80°C.

Expression and purification of PfCDPK1 and PfPKG recombinant proteins was conducted as described previously (*1*)

#### Kinase assay development

Enzyme kinase assays were conducted in 10µL reaction volume at room temperature for 1 hour in buffer containing 50 mM HEPES pH 7.5, 10 mM MgCl_2_, 1 mM EGTA, 2 mM DTT and 0.01% Tween20 in a black 384-well microplates (Greiner Bio-One). U*Light*™ substrates, Europium labelled anti-phospho-U*Light*™, and 10X Detection Buffer were purchased from Perkin Elmer. U*Light*-CREBtide (Sequence: CKRREILSRRPSYRK) and U*Light*-MBP peptide (sequence: CFFKNIVTPRTPPPSQGK) were the substrates of choice for *Pf*CLK1 and *Pf*CLK3, respectively. To determine K_m_ ATP, Serial dilutions of ATP from 1000 or 500 µM were added to the reaction wells containing *Pf*CLK1 + U*Light*-CREBtide or *Pf*CLK3 + U*Light*-MBP in 384 well black microplates (Greiner Bio-One). End point 665/615 nm data were fit to the Michaelis-Menten equation using GraFit version 5.0.12 (Erithacus Software Ltd.)

Reactions were performed using ATP at K_m_ concentration and optimized U*Light*™ and kinase concentrations (30 µM ATP and 5nM *Pf*CLK1 and 10µM ATP and *Pf*CLK3). Kinase reactions were stopped by addition of the Detection Mix Solution containing 10 mM EDTA and 1 mM anti-phospho-U*Light*™ in 1X Detection Buffer and the TR-FRET signal (emission1 = 665 nm, emission2 = 615 nm) was acquired after 1-8 hours using an EnVision™ Multilabel plate reader (Perkin Elmer, Waltham, MA).

Miniaturization to 1536-well format was possible and final assay performance was assessed by calculation of means and standard deviation of standard inhibition curves. Signal-to-background ratios S/B of 10 and 5 and Z´values of 0.8 and 0.6 were calculated for *Pf*CLK1 and *Pf*CLK3, respectively, suitable for HTS.

#### High-Throughput Screening

GSK compounds from the Tres Cantos Antimalarial Set (TCAMS, 13194 compounds and TopUp, 1500 compounds), and Protein Kinase Inhibitor Set displaying known anti-plasmodial activity (PKIS, 1115 compounds), and MRC-Technology index library (MRCT, 9970 compounds) were screened in single shot against *Pf*CLK1 and *Pf*CLK3. Kinase inhibitor-like compounds from the MMV Malaria Box (a selection of 260 from the original 400 compound library) were screened against *Pf*CLK1 and *Pf*CLK3 in dose response experiments (n=2), together with hits obtained from the Single Shot Screen.

Compounds were tested in single shot at 10 µM, or in dose response from 100 µM (11-point, 3-fold serial dilutions). Screening was performed in 1536-well plates, with final reaction and read-out volumes of 4 µL and 6 µL, respectively.

40 nL of 100x compound, or 40 nL DMSO (columns 11&12 - maximum and 35&36 - minimum activity controls) were dispensed onto each plate using a Echo Liquid handler (version?). 2µL of 2x PfCLK or kinase buffer were added onto all plates except columns 35 & 36 (no enzyme, minimum activity control), where 2 µL of kinase buffer were added. Reactions were initiated upon addition of 2x Substrate Solution containing K_m_ ATP concentration and U*Light*-MBP or U*Light*-CREBtide for PfCLK3 and PfCLK1, respectively.

Solutions were dispensed with a Multidrop Combi (Thermo Fisher Scientific, Waltham, MA, USA) and TR-FRET readouts were collected using an EnVision™ multilabel plate reader (Perkin Elmer) and normalized to control values of uninhibited and no enzyme.

The results from the High-Throughput screening were further analysed using Activity Base (ID Business Solutions Ltd., Surrey, UK). For each test compound, % inhibition was plotted against compound concentration. To calculate IC_50_, data was fit to the standard single-site four-parameter logistic equation:

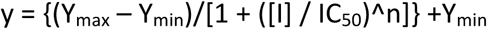

Where, Y_max_ is the maximum response, Y_min_ is the baseline, [I] is the concentration of compound, n is the Hill slope, IC_50_ is the inflection point concentration. In cases where the highest concentration tested (i.e., 100 μM) did not result in greater than 50% inhibition, the IC_50_ was determined as greater than 100 μM. pIC_50_= −logIC_50_, expressing the IC_50_ in molar units.

#### Parasite culture

*P. falciparum* cultures were maintained in RPMI-1640 media supplemented with 0.2% sodium bicarbonate, 0.5% Albumax II and 10mg/L gentamycin. Human blood group O, Rh +ve were used for parasite culture and cultures were maintained at 5% carbon dioxide, 5% oxygen and 90% nitrogen mixed gas incubator at 37°C.

#### Evolution of compound-resistant lines and whole genome sequencing

The *P. falciparum* Dd2 strain was cultured in triplicate in the presence of increasing concentrations of TCMDC-135051 to generate resistant mutants as previously described(*2*). After approximately 60 days of selection parasites were cloned in 96-well plates by limiting dilution(*3*). The half maximal (50%) inhibitory concentration was determined in dose-response format using a SYBR Green-I based cell proliferation assay as previously described(*4*). To determine genetic variants that arose during selection genomic DNA was isolated from each parasite strain. 250 base pair paired-end DNA libraries were prepared using the Illumina Nextera-XT method and sequenced on an Illumina Mi-seq according to manufacturer’s instructions. Reads were aligned to the *P. falciparum* 3D7 reference genome (PlasmoDB v9.0) as previously described(*5*). Single-nulceotide variants were detected using the Genome Analysis Toolkit (GATK v1.6) and the filtered based on the distribution of each quality control parameter (QD < 15.0, SQR > 1.9, MQ < 50.0, DP > 2700, MQRankSum −11.3 < x < 11.3, Qual > 400, DP > 6, GQ > 15.0, ReadPosRankSum −12.3 < x < 12.3, snp cluster: 3 in a window of 10.

#### Blood stage development inhibition assay by microscopy

Parasite cultures were tightly synchronized by two Percoll treatments in a span of one hour followed by sorbitol treatment to remove any left over schizont after second Percoll treatment and the very early ring stage parasites (0-1 Hour) were collected which were used to monitor inhibition of blood stage development of the parasite following treatment with inhibitor. Synchronized parasite cultures were treated with 1 μM TCMDC-135051 at time point 0, 10, 20, 30 and 40 hours and thin smears from each treatment or no treatment were prepared at time point 0 hour, 10 hour, 20 hour, 30 hour, 40 hour, 50 hour and 70 hour. Thin smears were stained with Giemsa stain and image of the parasites were acquired under light microscope at 100X magnification to study the stages of parasite blood stage development affected by the inhibition of *Pf*CLK3.

#### Asexual EC_50_ determination

Early ring stages of *P. falciparum* cultures were plated in 96-well black plate with clear bottom at 2 % hematocrite and 0.3 % parasitamia. Serial dilations of compound TCMDC-135051 staring at maximum 100 μM final concentrations were added to the parasite culture in triplicate and three wells were included with no drugs as control. Parasite cultures were incubated at 37°C for 72 hours and the viability of parasites in each condition were measured by staining parasite nuclei with nucleic acid stain SYBR Green (Invitrogen). Excitations at green channel were acquired on ClarioStar and the data were analyzed by Prism software.

#### Parasite reduction rate assay

Ring stage of 3D7 parasite strain was treated with TCMDC-135051 corresponding to 10X EC_50_ of the inhibitor concentration at blood stage development inhibition assay (EC_50_-188nM using hypoxanthine incorporation assay). Parasite cultures were treated for 120 hours with drug being renewed every 24 hour during the treatment. Samples of drug treated parasites were collected every 24-hour (24, 48, 72, 96 and 120 hour time points) and drug was washed from the collected sample by two rounds of centrifugation. Drug free parasites were diluted by three fold limiting dilution and cultured for 28 day in microtitre plate to allow all wells with viable parasites. Dilutions were performed four independent times to study the reproducibility of the experiment (n=4). Samples were collected on 21 day and 28 day from the microtitre plate to confirm parasite growth or no growth by hypoxanthine incorporation assay. For hypoxanthine incorporation assay 100μl of parasite cultures in a microtitre plate were supplemented with 8 μl of 0.025mci/ml ^3^H-hypoxanthine and parasite cultures were maintained for 72 hours and incorporation of ^3^H-hypoxanthine were measured by counting radioactivity in each well. Treatment with pyrimethamine was also conducted in parallel as control and pervious data for artemisnin, atovaquone and chloroquine were included to allow comparative classification of the killing profile.

#### Generation of G449 mutant parasite

Fragment of PfCLK3 gene containing part of exon2, exon 3, exon 4 and part of exon 5 (1143 bp) corresponding to 655-1797 bp in clk3 genomic sequence was amplified using primer CLK3-HR1 and CLK3-HR2 and the amplified product named as CLK3 Homologous region (CLK3-HR) was cloned in pHH1 derived vector using restriction sites HpaI and BglII. The rest of clk3 gene sequence down stream of CLK3-HR corresponding to 1798-3152 bp clk3 genomic sequence were modified by removing introns and the stop codon and the coding sequence was optimized for *E.coli* codon usage to make it dis-similar to clk3 genomic sequence. This fragment of gene which we named as CLK3-codon optimized (CLK3-CO) was commercially synthesized and included BglII recognition site at 5’ and XhoI recognition site at 3’. CLK3-CO was cloned downstream of CLK3-HR region in the parent plasmid using BglII and XhoI restriction sites in such a way that the triple HA tag sequence in the parent plasmid remained in frame with the clk3 sequence. The BglII restriction site that was artificially introduced for cloning purpose was mutated back to original CLK3 coding sequence by site directed mutagenesis using CLK3-BglII-KN1 and CLK3-BglII-KN2 primers. Site directed mutagenesis was used again to mutate Glycine in CLK3 at position 449 to an alanine residue using CLK3-G449P1 and CLK3-G449P2 primers. The target vector generated was sequenced to confirm the desired sequence of clk3 in the plasmid and further confirmed that 3-HA sequence is in frame with clk3 coding sequence. 10 μg of the target vector was used for transfection of schizont stage of the parasite using Lonza nucleofactor following the protocol as has been published by (*6*). Transfected parasites were cultured under selection drug pressure (2.5nM WR99210) till ring stage of the parasites could be seen in Giemsa stained smear. The parasites were maintained under drug OFF (three weeks) and drug ON (three weeks) pressure for two cycles to eliminate the parasite where the target plasmid was maintained episomally. Following drug cycle clones of parasite were generated by limiting dilution and the integration of target plasmid at the clk3 genomic locus of the cloned parasite was confirmed by PCR amplification of 1493 bp gene fragment using primers P3 (448-478 bp in clk3 genomic locus, upstream of CLK3-HR region) and P4 (1923-1940 bp sequence in clk3 gene of transgenic parasite, part of CLK3-CO region in target plasmid) which could be generated only if the target plasmid is integrated at the clk3 genomic locus. At the same time PCR amplification using primers P1 (571-605 bp in clk3 genomic locus, upstream of CLK3-HR region) and P2 (1825-1852 bp in clk3 genomic locus, in a region which is replaced by CLK3-CO region in transgenic parasite) did not show amplification in CLK3-G449P clones (A3 and A8), which confirms that the wild type clk3 locus was destroyed in the transgenic parasites. Amplification of 1282 bp product using primer P1 and P2 confirms that these primers can amplify clk3 at genomic locus of wild type 3D7 parasites.

#### Western blotting

Parasite lysate from 3D7 wild type parasite and two CLK3-G449P clones (A3 and A8) using lysis buffer containing 20 mM Tris pH-8.0, 150 mM sodium chloride, 1 mM di-thiotritol, 1 mM EDTA, 1% IGEPAL CA630 detergent and protease and phosphatase inhibitor mix. Parasite lysates were cleared by centrifugation at 21000xg for 5 minutes and 20 μg of parasite lysate were separated on 8% polyacrylamide gel under reducing condition. Proteins were transferred to nitrocellulose membrane and the membranes were blocked for two hour with blocking buffer that contain 20 mM Tris pH 7.4, 150mM sodium chloride, 0.1% Tween-20 and 5% skimmed milk. Following blocking the membranes were incubated with primary antibodies (Anti-HA monoclonal antibody or anti-*Pf*CDPK1 polyclonal antibody) at 1:1000 dilution in blocking buffer for one hour. The membranes were washed four times, each of 15 minutes, with wash buffer (blocking buffer without milk) and incubated with goat anti-rat HRP conjugated antibody at 1:2000 dilution in blocking buffer for one hour. Following this incubation the blots were washed four times with wash buffer and developed using enhanced chemi-luminiscence kit. The luminescences from the blots were acquired on X-ray film. The control gel following electrophoresis was stained with Coomassie stain to show equal loading.

#### DNA Microarray experiment

Microarray was used to study changes in transcript level in the parasite following treatment with CLK3 inhibitor. Trophozoite stage of wild type (Dd2) or mutant (051C) parasites were purified using MACS (magnetic assisted cell sorting, Milteny Biotec) purification column and treated either with compound TCMDC135051 (1μM) or solvent DMSO for one hour at 37°C. Following inhibitor treatment the parasites were treated with 0.1% saponin in phosphate buffer saline (PBS) for one minute on ice to release parasites from host erythrocyte. Parasites were washed twice with PBS and the parasite pellets were used for total RNA isolation using RNeasy Lipid Tissue Kit (Qiagen). Bioanalyzer 2100 instrument (Agilent) was used to assess the integrity of RNA samples and samples with RNA integrity score of >8 were used for the array experiment. cDNA was prepared from the RNA samples using cDNA synthesis kit (Agilent) and equal amounts of cDNAs (amount 200ng) were labeled with Cy3 label and the equal incorporation of label was confirmed by NanoDrop. Custom DNA microarray slides were prepared (Agilent) that contained 14353 unique probes covering 5752 P. falciparum genes as described before by(*7*). Microarray slides were hybridized with labeled cDNA samples and intensity of label in each probe of the array were acquired by Agilent G2505C Microarray scanner. Normalized data from four independent biological replicates were used to identify probes showing at least two-fold change between the control samples and inhibitor treated samples. Statical significance (pValue 0.5<) between the control and treated samples were calculated using moderate T-test analysis.

#### Liver stage development assay

*Plasmodium berghei* sporozoites expressing Luciferase were freshly collected from salivary gland of *Anopheles stephensi* mosquitos to be used for measurement of inhibition of exoerythrocytic form (EEF) development by drug treatment. The freshly collected sporozoites were used for infection of human hepatocarcinoma HepG2 cells expressing the tetraspanin CD81 receptor (HepG2-A16-CD81EGFP cells). HepG2 cells were pretreated for 18 hours with the inhibitor before infection with *P. berghei* sporozoites. After infection, the HepG2 cells were further treated with the drug at 12-point serial dilution for 48 hours and the inhibition of EEF development were measured by bioluminescence. Atovaquone and DMSO treatments were used as positive and negative controls in the microtitre based plate assay. IC50 value of the inhibitor for inhibition of sporozoite development were generated using bioluminescence values.

#### *P. berghei* blood stage development assay

*P. berghei* ANKA strain of parasite was used for infection of Balb/c mice. Blood of *P. beghei* infected mice were collected at parasitemia of 1-3% and the parasites were allowed to grow in *in vitro* for 24 hour, which allowed the parasites to synchronize at late schizont stage. Mice were infected with late schizont *P. berghei* parasites for 2-4 hours that allowed schizonts to rupture and form nascent ring stage of the parasite. Ring stage parasite infected RBCs were collected from the mice and treated with drug at eight-point dilution for 22-24 hours at 37 ^0^C in a microtitre plate. Follwing inhibitor treatment, parasites were stained with Hoechst DNA stain and acquired by flow cytometry to measure percentage of schizont-infected cells. Artemisinin and DMSO treatments were used as positive and negative control. Growth inhibitory concentrations were calculated using Prism 7 software.

#### Stage II gametocyte inhibition assay

*P. falciparum* Pf2004 strain of parasites expressing tdTom as marker for stage II gametocyte were used to measure inhibition of gametocyte development by the inhibitor, as previously described(*8*). At 28 hours post invasion (28hpi) the regular culture medium was replaced with serum-free medium (-SerM) complemented with a minimal fatty acid mixture and bovine serum albumin (0.39% fatty acid-free BSA and oleic and palmitic acid [30 μM each; added from 30 mM ethanol-solved stocks]) to induce sexual commitment. After 20 hours of incubation in −SerM the medium was replaced with complete medium containing 10% serum for the remainder of the experiment. Inhibitor at eight-point dilutions starting at 20μM was added at 28hpi, and it was maintained for 42 hours to study the effect of the inhibitor on early gametocyte (stage II) development. Following treatment gametocytemia was measured by flow cytometry using tdTom reporter fluorescence as a viability marker (details on cytometry are given in(*8*). The inhibitory effect of the compound on gametocyte development was calculated using Graph Prism 7 software.

### DNA primers used

**CLK1-FamB1**

AGGAGATATACATATGGATGATGAAATTGTCCATTTTAGTTGG

**CLK1-FamB2** GAAGTACAGGTTCTCTTCAAGGAACTTGTGCTTTAATAATTCG

**CLK3-HR1**

ATGC**GTTAAC**GAAATACCATCTAATCCATCATATATCGACC

**CLK3-HR2**

ATGC**AGATCT**ATATTTTATACTACTTAATAAACGAATGATGTGCCT

**CLK3_BglII-KN1**

Cattcgaacactaagcataaatgatttttatattttatactacttaataaacgaatgatgtgccttttattgtcc

**CLK3_BglII-KN2**

Ggacaataaaaggcacatcattcgtttattaagtagtataaaatataaaaatcatttatgcttagtgttcgaatg

**CLK3-G449P1**

Gggcaatacgcagattgggccacatccattcgaaca

**CLK3-G449P2**

Tgttcgaatggatgtggcccaatctgcgtattgccc

**P1**

*GTATTTTCAGGGCGCC*Cagaatgatcagataaataaagattataacaatga

**P2**

GTGCTATTCTTAAGTTACCCCACATCCA

**P3**

CGAACACAAGAAAATGAGGATAAACTTCTAG

**P4**

CGCATGTGACGCAGGGCA

## Supplementary Figures

**Supplementary Figure S1.**
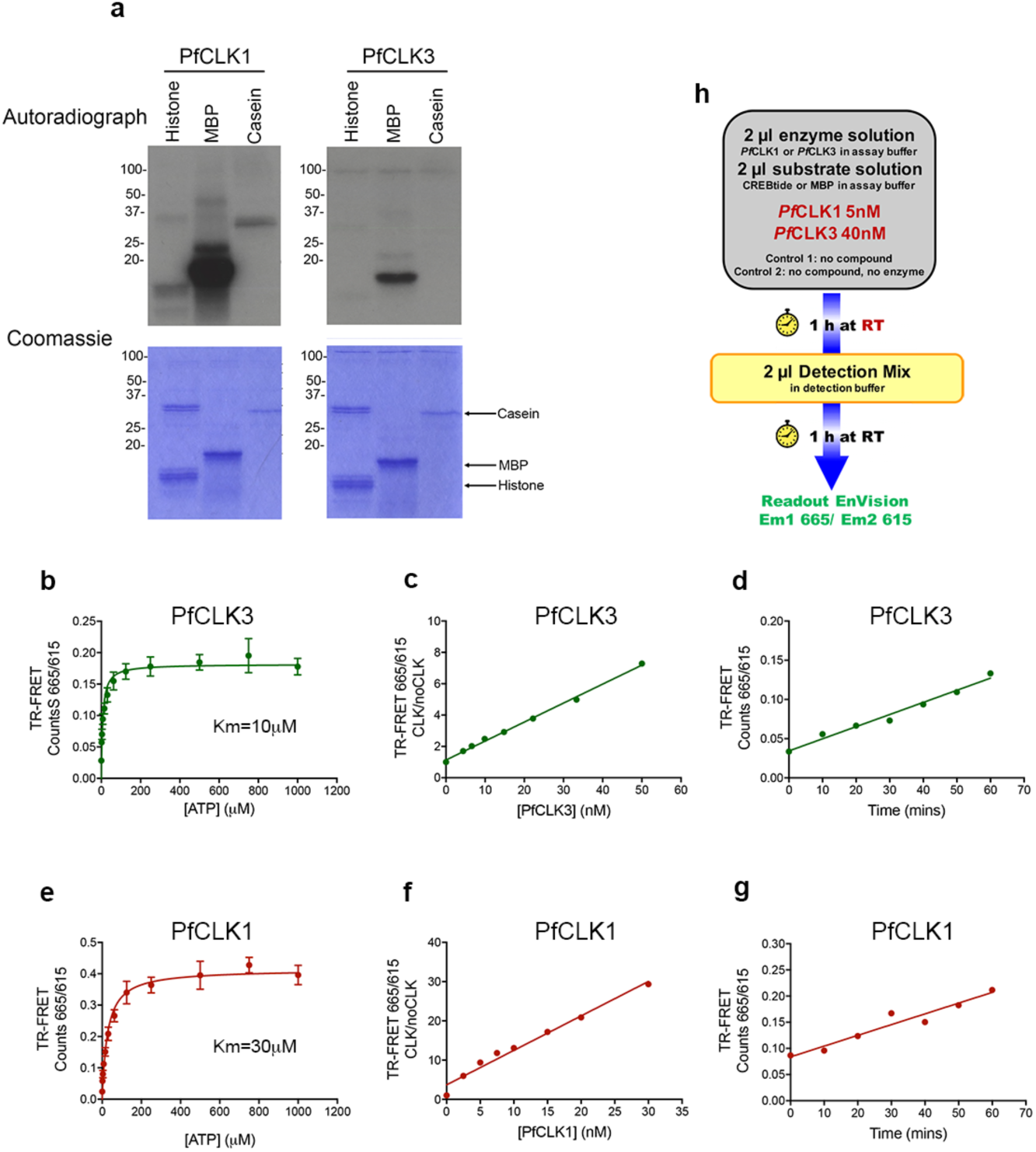
Generation of a HTS screen for *Pf*CLK1 and *Pf*CLK3. **(a)** Gel based assay of the phosphorylation of myelin basic protein (MBP) *α*-casein and histone by recombinant *Pf*CLK1 and *Pf*CLK3. The top gel is an autoradiograph and the bottom a Commassie stain of the same gel as a loading control. The position of MBP, *α*-casein and histone are shown. **(b)** Measure of Vmax and Km for ATP of recombinant *Pf*CLK3 **(c)** Linear relationship between the activity read out of the TR-FRET kinase assay and the concentration of *Pf*CLK3 using in the kinase reaction. **(d)** Time course of the TR-FRET kinase reaction for *Pf*CLK1 when used at an enzyme concentration of 40nM. **(e)** Measure of Vmax and Km for ATP of recombinant *Pf*CLK1 **(f)** Linear relationship between the activity read out of the TR-FRET kinase assay and the concentration of *Pf*CLK1 using in the kinase reaction. **(g)** Time course of the TR-FRET kinase reaction for *Pf*CLK1 when used at an enzyme concentration of 40nM. **(h)** Scheme of the assay protocol for *Pf*CLK1 and *Pf*CLK3 high-throughput screens. The data shown in the graphs are the mean ±S.E.M of at least three independent experiments.

**Supplementary Figure S2.**
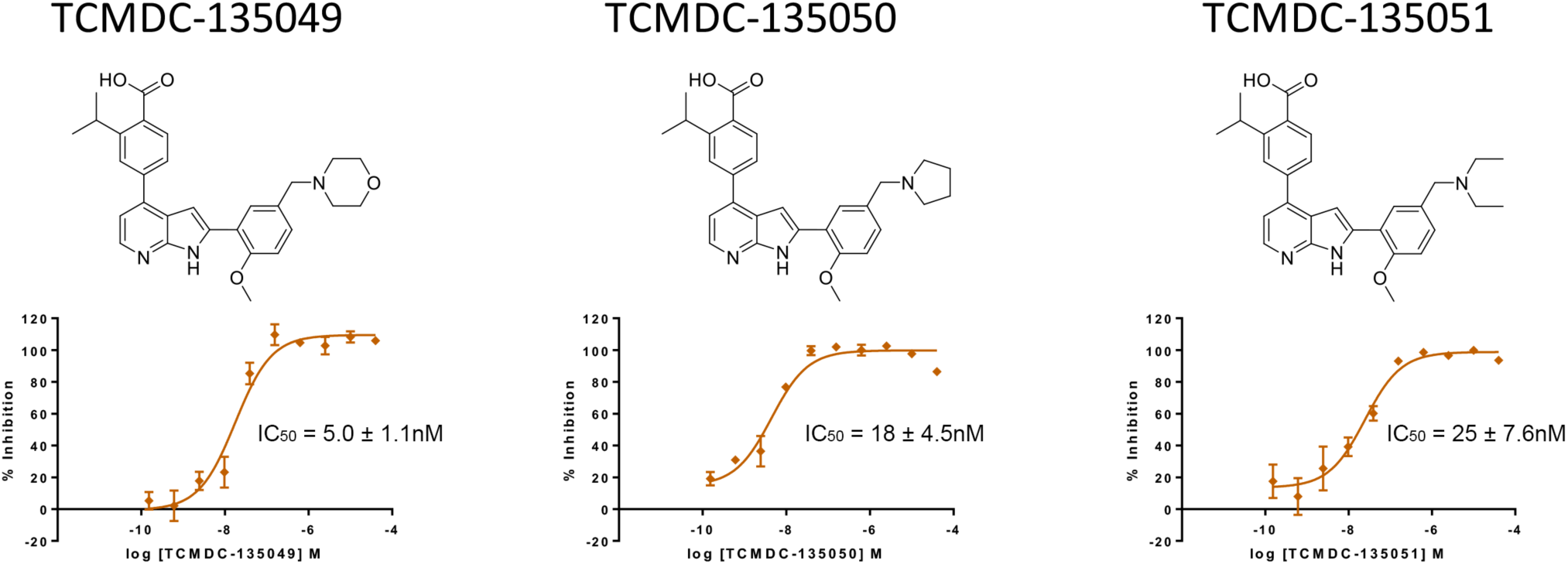
Series of related molecules identified from HTS as *Pf*CLK3 inhibitors. TR-FRET assay of *Pf*CLK3 activity was used to generate concentration inhibition curves of TCMDC-135049, TCMDC-135050 and TCMDC-135051. Shown is the mean of three experiments ± S.E.M).

**Supplementary Figure S3.**
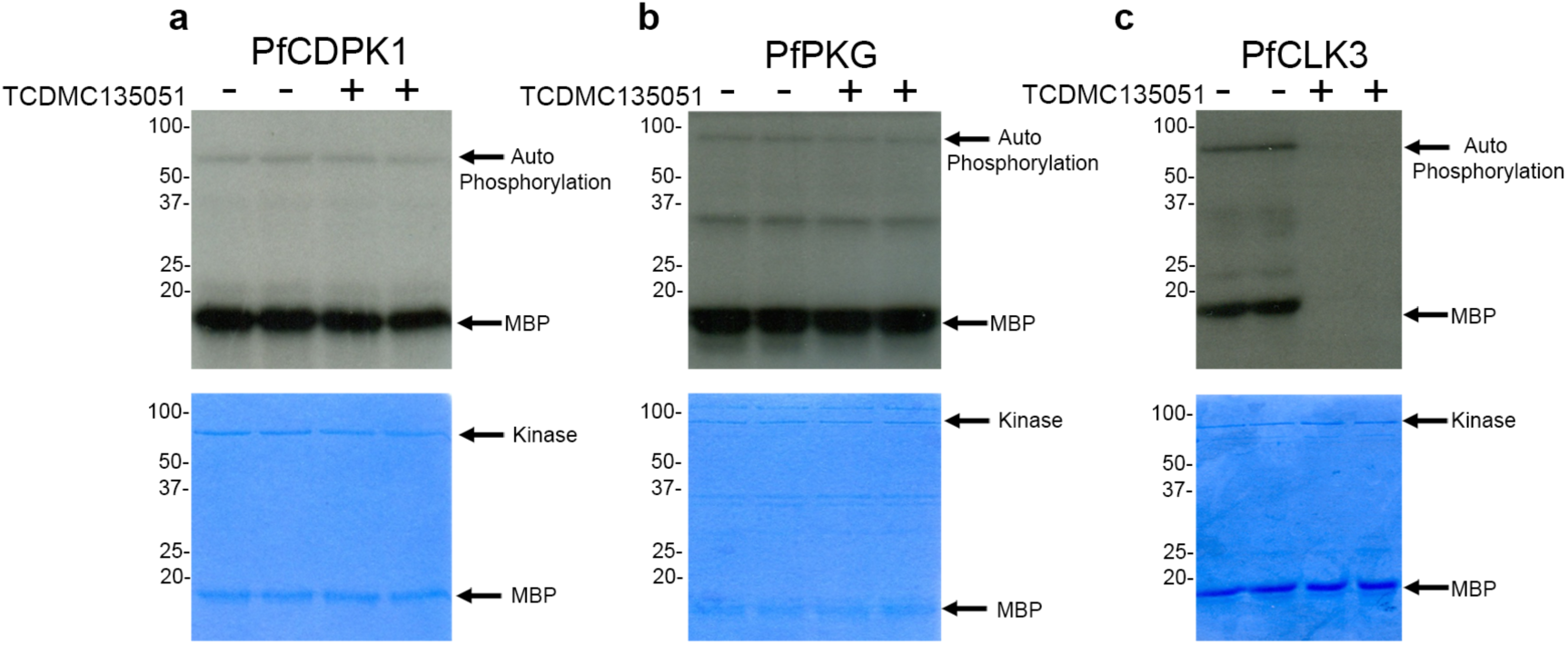
TCMDC-135051 activity against P*f*CDPK1 and *Pf*PKG. Gel based assay of the phosphorylation of myelin basic protein (MBP) by **(a)** *Pf*CDPK1, **(b)** *Pf*PKG and **(c)** *Pf*CLK3. The top gel is an autoradiograph and the bottom a Commassie stain of the same gel as a loading control. The position of MBP and auto-phosphoryated kinase and the recombinant kinase is shown.

**Supplementary Figure S4.**
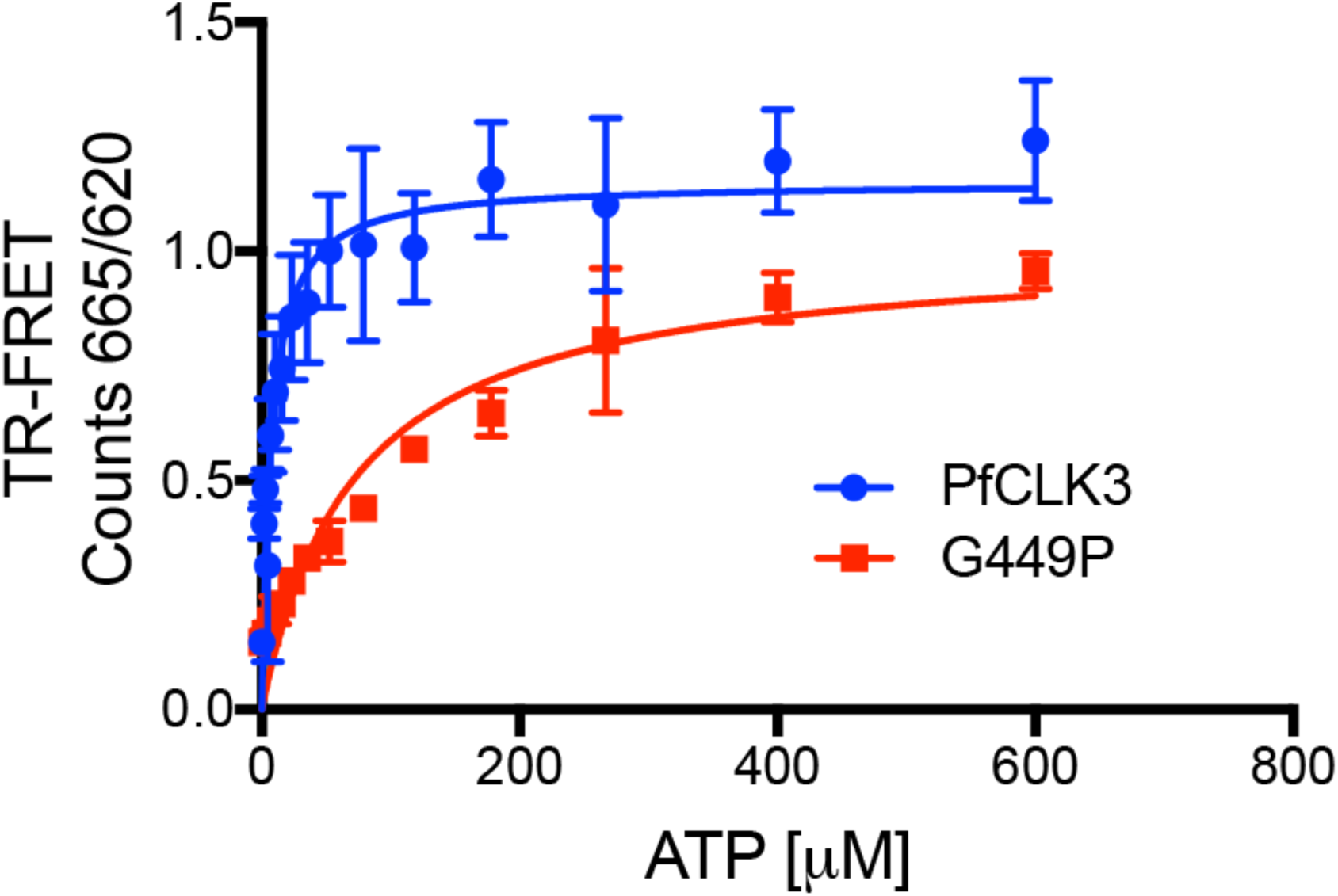
Comparison of the enzyme kinetics of *Pf*CLK3 and the G449P variant. Measure of recombinant *Pf*CLK3 and G449P at various concentrations of ATP. The data shown is the mean ±S.E.M of at least three independent experiments.

**Supplementary Figure S5.**
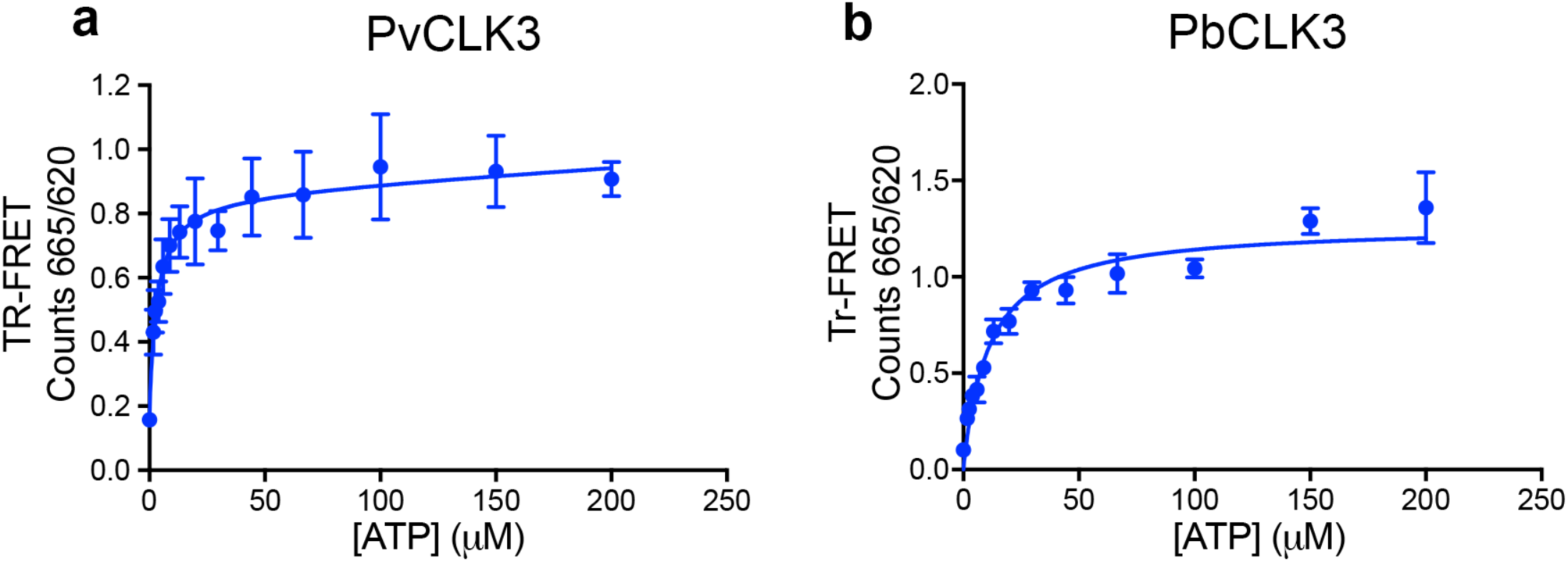
Comparison of the enzyme kinetics of *Pv*CLK3 and *Pb*CLK3. Measure of recombinant **(a)** *Pf*CLK3 and **(b)** *Pb*CLK3 at various concentrations of ATP. The data shown is the mean ±S.E.M of at least three independent experiments.

**Supplementary Figure S6.**
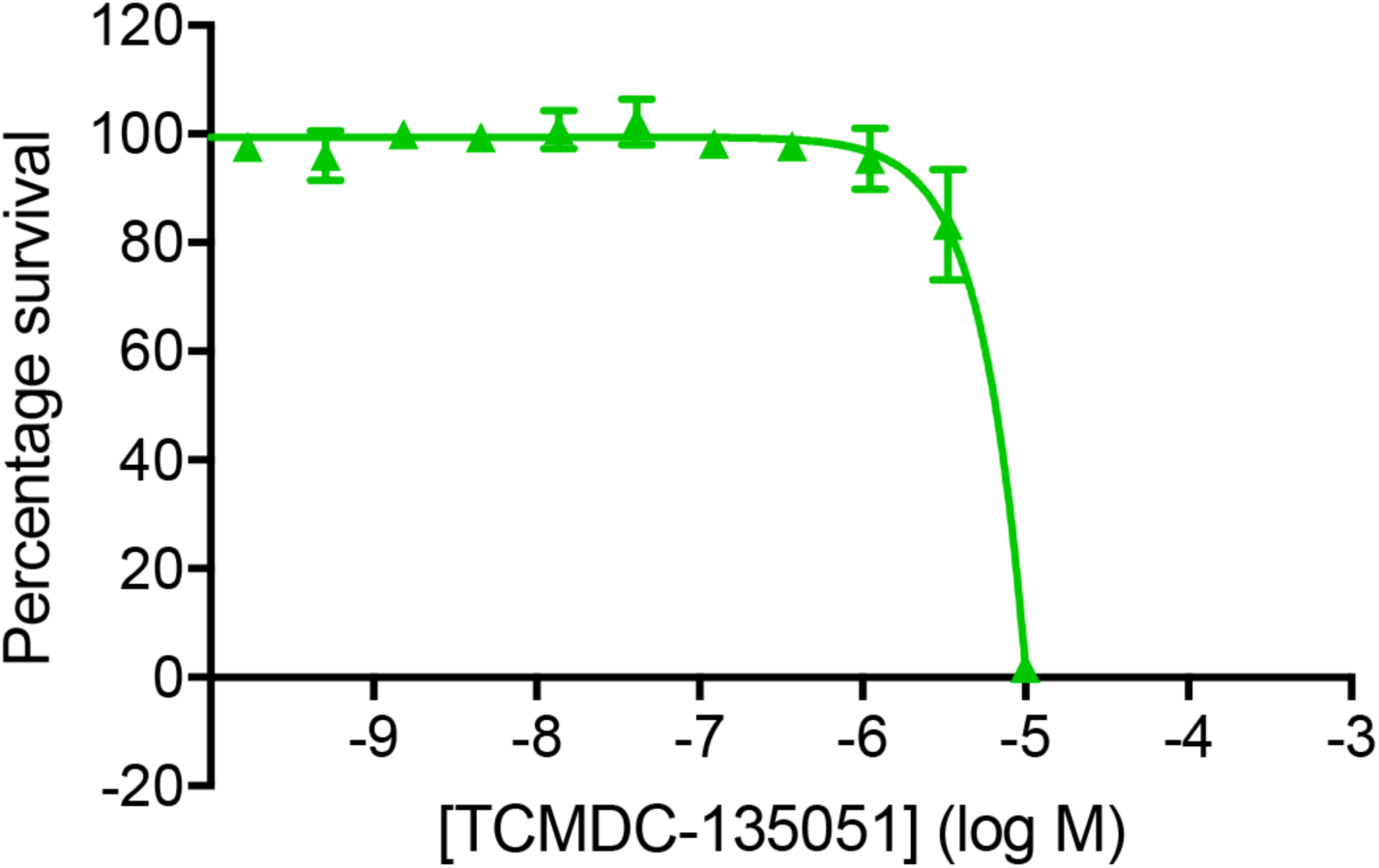
HepG2 viability assay. HepG2 cells were treated with the indicated concentrations of TCMDC-13501 for 48 hours after which cell viability was determined by a bioluminescence assay. The data shown is the mean ±S.E.M of at least three independent experiments.

## Supplementary Tables

**Supplementary Table S1. Compound Screen.**

Hits from the primary single dose HTS (2838) and compounds contained in the MMV-malaria box (259) were used in concentration inhibitor experiments. Shown are pIC50 values against PfCLK3, PfCLK1, PvCLK3 and hCLK2 (sheet 1). These data are divided into data associated with TCAMs/GSK compounds (sheet 2), PKIS compounds (sheet 3), MRCT compounds (sheet 4) and MMV compounds (sheet 5).

**Supplementary Table S2. Transcription changes in parent *P. falciparum* line Dd2 following treatment with TCMDC-135051.**

*Sheet 1* – Statistically significant changes (t-test n=4) in transcript levels following treatment of Dd2 parasites at trophozoite stage with TCMDC-135051 (1μM for 60 mins). Shown are the probes used and changes associated with each of the probes. *Sheet 2* – Summary of the probes that reveal significantly down-regulated genes following TCMDC-135051 treatment. *Sheet 3* - Summary of the genes that are significantly down-regulated following TCMDC-135051 treatment. *Sheet 4* - Summary of the probes that reveal significantly up-regulated genes following TCMDC-135051 treatment. *Sheet 5*-Summary of the genes that are significantly up-regulated following TCMDC-135051 treatment.

**Supplementary Table S3. Transcription changes in the resistant parasite line TM051C following treatment with TCMDC-135051.**

*Sheet 1* – Statistically significant changes (t-test n=4) in transcript levels following treatment of TM051C parasites at trophozoite stage with TCMDC-135051 (1μM for 60 mins). Shown are the probes used and changes associated with each of the probes. *Sheet 2* – Summary of the probes that reveal significantly down-regulated genes following TCMDC-135051 treatment. *Sheet 3* - Summary of the genes that are significantly down-regulated following TCMDC-135051 treatment. *Sheet 4* - Summary of the probes that reveal significantly up-regulated genes following TCMDC-135051 treatment. *Sheet 5*-Summary of the genes that are significantly up-regulated following TCMDC-135051 treatment.

**Supplementary Table S4. Transcription changes designated as a result of on-target inhibition of *Pf*CLK3 and a comparison with essential *Plasmodium falciparum* genes.**

Subtracting the genes where transcription is significantly changed following treatment with TCMDC-135051 (1μM for 60 mins) in TM051C parasites with those changed in Dd2 parasites revealed the “on target” transcriptional changes associated with inhibition of PfCLK3. *Sheet 1* – On target up-regulated genes. *Sheet–2* Identification (highlighted) of the on target up regulated genes that are essential for parasite blood stage survival. *Sheet 3-* On target down-regulated genes. *Sheet 4-* Identification (highlighted) of the on target down regulated genes that are essential for parasite blood stage survival. Also shown are the allocation of gene function to those down regulated genes.

## Acknowledgements

This work has been funded through an MRC Toxicology Unit programme grant (ABT, MMA), Lord Kelvin Adam Smith Fellowship (MMA), GSK Open Lab Foundation Award (ASA), joint MRC Toxicology Unit and MRC Unit the Gambia PhD programme (OJ), Daphne Jackson Fellowship (DM). EAW is supported by grants from the NIH (5R01AI090141 and R01AI103058) and by grants from the Bill & Melinda Gates Foundation (OPP1086217, OPP1141300) as well as by Medicines for Malaria Venture (MMV). Drug WR99210 for selection of transgenic parasites were gifted by Jacobus Pharmaceuticals. Our thanks to Emma Barr for supporting the work of the Tobin group.

